# The Most Parsimonious Reconciliation Problem in the Presence of Incomplete Lineage Sorting and Hybridization is NP-Hard

**DOI:** 10.1101/2021.03.14.435321

**Authors:** Matthew LeMay, Yi-Chieh Wu, Ran Libeskind-Hadas

## Abstract

The maximum parsimony phylogenetic reconciliation problem seeks to explain incongruity between a gene phylogeny and a species phylogeny with respect to a set of evolutionary events. While the reconciliation problem is well-studied for species and gene trees subject to events such as duplication, transfer, loss, and deep coalescence, recent work has examined species phylogenies that incorporate hybridization and are thus represented by networks rather than trees. In this paper, we show that the problem of computing a maximum parsimony reconciliation for a gene tree and species network is NP-hard even when only considering deep coalescence. This result suggests that future work on maximum parsimony reconciliation for species networks should explore approximation algorithms and heuristics.

## 1 Introduction

Genes evolve via several evolutionary processes operating at various evolutionary timescales. Nucleotides can mutate, and domains can recombine. Genes can be generated, lost, or replaced through gene duplication, gene loss, horizontal gene transfer, and gene conversion. Populations can diverge or combine through speciation and hybridization. In addition to these events, within a population, polymorphisms can persist across speciation events, leading to a phenomenon known as incomplete lineage sorting. Thus, the history of a set of genes may differ from the history of the species in which they evolved.

In phylogenetics, *reconciliations* attempt to explain these differences by mapping gene histories within species histories to infer the evolutionary events that shaped that gene family. The simplest and most common approach seeks a most parsimonious reconciliation (MPR) (Goodman et al. 1979, Page 1994, Maddison 1997), in which each type of event in the model has an associated non-negative cost and the objective is to find a reconciliation of minimum total cost.

The time complexity of the MPR problem depends on the events being modeled and the set of constraints being considered. For example, the lowest common ancestor mapping, which can be computed in polynomial time, solves the MPR problem when considering only duplications (Górecki and Tiuryn 2006), only duplications and losses (Górecki and Tiuryn 2006), and only deep coalescence (Wu and Zhang 2011). When considering duplications, transfers, and losses, the MPR problem can be solved in polynomial time or is NP-hard depending on whether the species tree is undated, partially dated, or fully dated, and on whether the reconciliation is constrained to be time-consistent (Ovadia et al. 2011, Tofigh et al. 2011). Similarly, depending on details of the underlying model, the MPR problem can be solved in polynomial time when considering duplications, transfers, losses, and coalescence (Stolzer et al. 2012, Chan et al. 2017), or is NP-hard when considering duplications, losses, and coalescence (Bork et al. 2017). The MPR problem is also NP-hard when simultaneously modeling the evolution of domains, genes, and species (Li and Bansal 2019).

Though the species history is often represented by a tree, several authors have recently considered reconciliations with species networks which model species hybridization. For example, several methods exist for the related problem of inferring a species network that minimizes deep coalescence (Yu et al. 2011; 2013a;b). However, the authors did not analyze the time complexity of their algorithms. Furthermore, their approaches require searching over the space of species network topologies, in contrast to the problem considered here, in which the species network is assumed to be known. Other authors have focused on the problem of inferring MPRs between gene trees and species networks, showing that the problem can be solved in polynomial time when minimizing the duplication-transfer-loss cost (Libeskind-Hadas and Charleston 2009) or the duplication-loss cost (To and Scornavacca 2015). There is little previous work on the problem of inferring MPRs between a gene tree and species network in which incongruence is due to deep coalescence, and the question of whether this problem can be solved in polynomial time remained open.

In this paper, we show that the problem of inferring an MPR between a gene tree and species network in the presence of incomplete lineage sorting is NP-hard. Our results suggests that future work on this problem should focus on developing heuristics or approximation algorithms.

## 2 Definitions

We use the terms *node* and *vertex* interchangeably. A *rooted phylogenetic network* refers to a rooted directed acyclic graph with a single root with in-degree 0 and out-degree 2; additional internal nodes with either in-degree 1 and out-degree 2, called *branch nodes*, or in-degree 2 and out-degree 1, called *hybridization nodes*; and one or more leaves with in-degree 1 and out-degree 0,. Edges leading to hybridization nodes are called *hybridization edges*. Given a network *N*, let *V* (*N*) denote its node set and *E*(*N*) denote its edge set. Let *L* (*N*) *⊂V* (*N*) denote its leaf set, *I*(*N*) = *V* (*N*) \ *L*(*N*) denote its set of internal nodes, and *r*(*N*) *∈ I*(*N*) denote its root node. For a node *v ∈ V* (*N*), let *c*(*v*) denote its set of children (the empty set if *v* is a leaf), let *p*(*v*) denote its set of parents (the empty set if *v* is the root node), and, if *v* has a single parent, *e*(*v*) denotes the edge (*p*(*v*), *v*). The size of *N*, denoted by |*N* |, is equal to |*V* (*N*)| + |*E*(*N*)|.

Let ≤ _*N*_ (*<*_*N*_) be the partial order on *V* (*N*) such that *v ≤*_*N*_ *u* (*v <*_*N*_ *u*) if and only if there exists a path in *N* from *u* to *v* (*v ≠ u*); *v* is said to be *lower or equal to* (lower than) *u*, and *v* a (strict) *descendant* of *u*, and *u* a (strict) *ancestor* of *v*. The partial order *≥* _*N*_ (*>*_*N*_) is defined analogously.

Given two nodes *u* and *v* of *N* such that *v ≤*_*N*_ *u*, a path from *u* to *v* in *N* is a sequence of contiguous edges from *u* to *v* in *N* .Note that if *u* = *v*, the path from *u* to *v* is empty. As there can be multiple paths between pairs of vertices in a network, let *paths*_*N*_ (*u, v*) denote the set of all paths from *u* to *v*. Let *paths*(*N*) denote the set of all paths in network *N*.

A phylogenetic tree is a phylogenetic network with no hybridization nodes; that is, a directed binary tree. In the remainder of this paper, we refer to rooted phylogenetic networks and rooted phylogenetic trees simply as *networks* and *trees*, respectively.

A *species network S* represents the evolutionary history of a set of species and a *gene tree G* represents the evolutionary history of a set of genes sampled from these species. A *leaf map Le*: *L*(*G*) *→L*(*S*) associates each leaf in the gene tree with a corresponding species from which the gene was sampled. The map need not be one-to-one nor onto. Note that gene phylogenies are assumed to be trees whereas species phylogenies may, in general, be networks.

### 2.1 Reconciliations

A *reconciliation* for a given gene tree, species network, and leaf map comprises a pair of maps: The *vertex map* associates each node of *G* with a node of *S*. The *path map* associates a path in *S* from the node of *S* where *p*(*v*) is mapped by the vertex map to the node of *S* where *v* is mapped by the vertex map. The vertex map must be consistent with the given leaf map and must satisfy the the temporal constraints of the species map; namely if a gene node *g* is mapped to species node *s* and a child *g*^*’*^ of *g* is mapped to species node *s*^*’*^, then *s* must be an ancestor of *s*^*’*^. The path map is required because *S* is a network and thus there may be multiple paths between ancestors and descendants in the network. The formal definition of reconciliations is given in Definition 2.1.

#### Definition 2.1

(Reconciliation). Given a gene tree *G*, a species network *S*, and a leaf map *Le*, a *reconciliation*^1^ *R* for (*G, S, Le*) is a pair of maps (*R*_*v*_, *R*_*p*_) where *R*_*v*_: *V* (*G*) *→ V* (*S*) is a *vertex map* and *R*_*p*_: *V* (*G*) *→ paths*(*S*) is a *path map* subject to the following constraints:

1. If *g ∈ L*(*G*), then *R*_*v*_(*g*) = *Le*(*g*).
2. If *g ∈ I*(*G*), then for each *g*^*’*^ *∈ c*(*g*), *R*_*v*_(*g*^*’*^) *≤*_*S*_ *R*_*v*_(*g*).
3. If *g ≠ r*(*G*), then *R*_*p*_(*g*) *∈ paths*_*S*_(*R*_*v*_(*p*(*g*)), *R*_*v*_(*g*)). Otherwise, *R*_*p*_(*g*) = ∅.

Constraint 1 asserts that *R*_*v*_ extends the leaf map *Le*. Constraint 2 asserts that *R*_*v*_ satisfies the temporal constraints implied by *S*. Constraint 3 asserts that the vertex map and path map are consistent.

In a multispecies coalescent process, evolution in the species network is viewed backward in time, from the leaves toward the root. Then, given a reconciliation *R*, we can count the number of gene lineages “passing through” each edge *e* of the species network. Specifically, given edge *e ∈ E*(*S*),

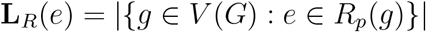

The number of “extra lineages” is defined to be:

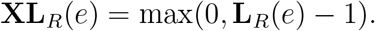

Note, for example, that if two gene paths pass through a species edge, there is one extra lineage on that edge.

Finally, the *deep coalescence cost* of a reconciliation is the sum of extra lineages across all edges of the species network:

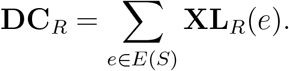

This value is the *reconciliation cost* in this model. Finally, we formalize these optimization and decision problems:

**Problem 2.1** (Most Parsimonious Reconciliation (MPR)). Given a gene tree *G*, a species network *S*, and a leaf map *Le*, find a reconciliation *R* for (*G, S, Le*) such that the deep coalescence cost **DC**_*R*_ is minimized.

**Problem 2.2** (Most Parsimonious Reconciliation Decision Problem (MPRD)). Given a gene tree *G*, a species network *S*, a leaf map *Le*, and an integer *k*, is there a reconciliation *R* for (*G, S, Le*) such that **DC**_*R*_ *≤ k*?

## 3 NP-hardness

### Theorem 3.1.

*MPRD is NP-hard*.

In the proof that follows, it will be convenient to consider the gene tree as the collection of all paths *P* from its root to its leaves. For a given leaf map *Le*: *L*(*G*) *→ L*(*S*), a *lineage map* with respect to *Le* is a mapping *M* from each path *p*_*𝓁*_ *∈ P* whose endpoint is leaf *𝓁* to a path in *S* from *r*(*S*) to *Le*(*𝓁*). Each reconciliation has a corresponding lineage map,*M*. Specifically, let *R* = (*R*_*v*_, *R*_*p*_) be a reconciliation and let *r*(*G*), *g*_1_, …, *g*_*k*_, *g*_*k*_ = *𝓁*, denote the nodes on the unique path from *r*(*G*) to leaf *𝓁*. Then, *M* associates this path in *G* with path *R*_*p*_(*g*_1_)*R*_*p*_(*g*_2_) … *R*_*p*_(*g*_*k*_) in *S*. Note that multiple different reconciliations may induce the same lineage map since there are, in general, different mappings of nodes of *G* to nodes of *S* than induce the same set of paths. For simplicity, when referring to lineage maps we use the notation *M* (*𝓁*) in lieu of *M* (*p*_*𝓁*_) where *p* is the unique path from *r*(*G*) to leaf *𝓁*.

Finally, we use the notation (*A, B*) to represent a binary tree with a root and children *A* and *B*, either of which may be leaves or trees themselves.

*Proof*. Our proof is by a reduction from 3SAT. Consider an instance of 3SAT with *n* variables and *m* clauses and, without loss of generality, assume that *n* and *m* are both powers of 2.

(If not, the 3SAT instance can be trivially padded with dummy variables and clauses to construct an equivalent instance in polynomial time.)

### Construction

The gene tree *G* is constructed as follows: A *gene variable gadget* for a variable *x*_*i*_ comprises a tree 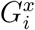 with three leaves, labelled *a*_*i*_, *b*_*i*_, *y*_*i*_, with the topology ((*a*_*i*_, *b*_*i*_), *y*_*i*_). The root of this gadget is labelled *x*_*i*_. These *n* variable gadgets are connected via a perfect binary tree *G*^*x*^. A *gene clause gadget* for a clause *c*_*j*_ consists simply of a single leaf *𝓁*_*j*_. These *m* leaves are connected via a perfect binary tree *G*^*c*^. A *k-comb* of length *k* is a binary tree constructed from a path of length *k* (*k* +1 vertices and *k* edges) where each of the first *k* vertices on that path has two children; one is the next vertex on the path and another is a leaf. In total a *k*-comb gadget then has *k* + 1 leaves. The root of the gene tree *G* has two children, each of which is the root of a 2*n*-comb, one called comb *d* and the other comb *e*. One of the two deepest leaves of comb *d* is the root of tree *G*^*x*^ and the remaining leaves are labeled *d*_1_, …, *d*_2*n*_ in order of depth from the root. Similarly, one of the two deepest leaves of comb *e* is the root of *G*^*c*^ and the remaining leaves are labeled *e*_1_, …, *e*_2*n*_ in order of depth from the root. The structure of the gene tree is depicted in Figure 1.

**Figure 1.**
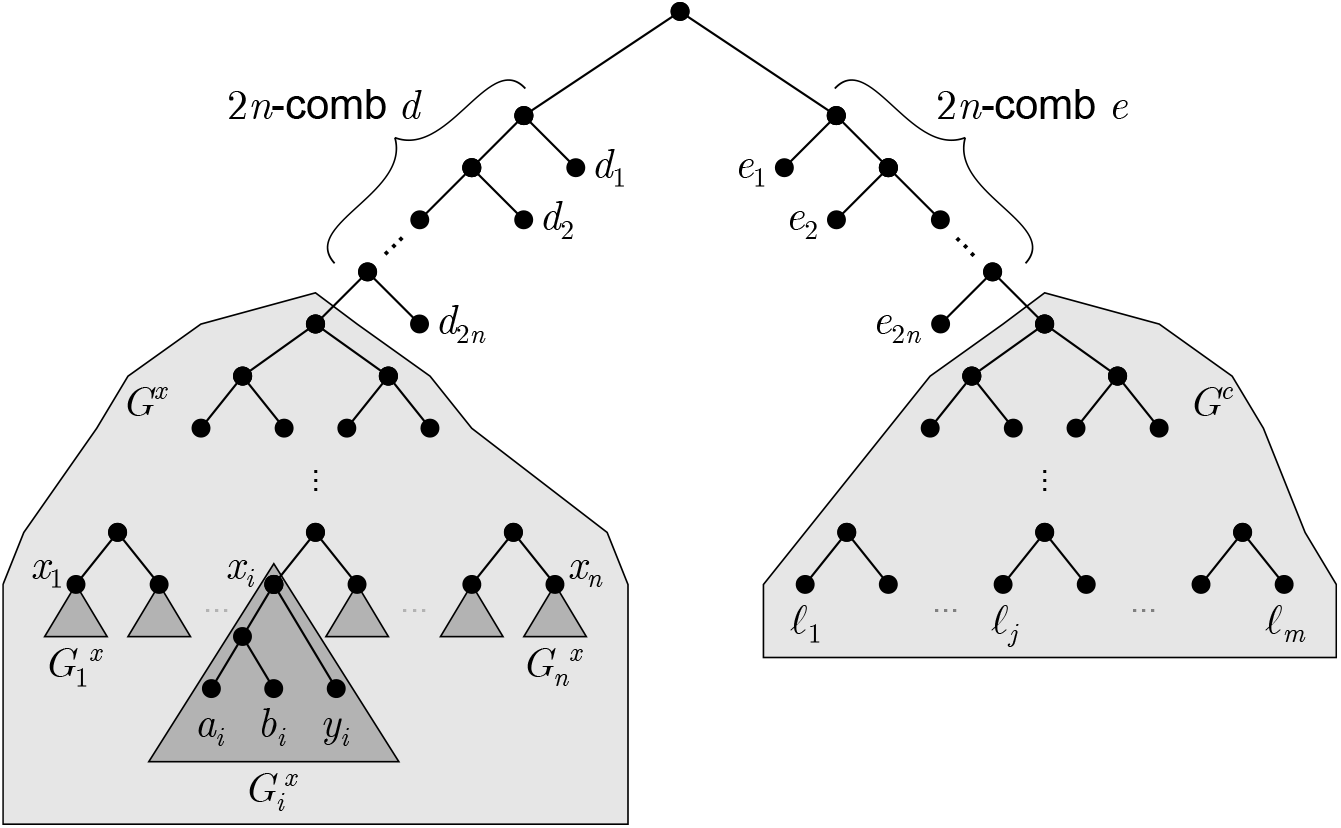
The gene tree in the NP-hardness construction. The shaded subtree *G*^*x*^ on the left contains a variable gadget 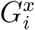 for each variable *x*_*i*_ in the 3SAT instance, and the shaded subtree *G*^*c*^ on the right contains a single leaf *𝓁*_*j*_ for each clause *j* in the 3SAT instance.

The species network *S* is constructed as follows: A *species variable gadget* for variable *x*_*i*_ consists of a subtree 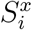 with four children labeled *T*_*i*_, *A*_*i*_, *B*_*i*_, and *F*_*i*_, with topology ((*T*_*i*_, *A*_*i*_), (*F*_*i*_, *B*_*i*_)). *A*_*i*_ and *B*_*i*_ are leaves. The root of this gadget is labelled *X*_*i*_. *T*_*i*_ and *F*_*i*_ are the first vertices of paths which correspond to setting *x*_*i*_ to true or false, respectively. We henceforth refer to these paths as the *variable setting paths* for *T*_*i*_ and *F*_*i*_, respectively. The remaining vertices on these variable settings paths are described in the next paragraph; ultimately these paths join at a hybridization node which has a single leaf child *Y*_*i*_. The roots of these *n* variable gadgets are joined via a perfect binary tree *S*^*x*^.

A *species clause gadget* 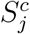 for clause *C*_*j*_ is constructed as follows. Let *z*_1_, *z*_2_, and *z*_3_ denote the three literals in that clause. If literal *z*_1_ is the unnegated variable *x*_*i*_ then a vertex *U*_1,*j*_ and its child *V*_1,*j*_ are introduced on the variable setting path for *F*_*i*_ in the species variable gadget for *x*_*i*_. Conversely, if literal *z*_1_ is the negation of *x*_*i*_, then vertex *U*_1,*j*_ and its child *V*_1,*j*_ are introduced on the variable setting path for *T*_*i*_. The analogous process is used to introduce a pair of vertices *U*_2,*j*_ and *V*_2,*j*_ for the variable setting path for literal *z*_2_ and a pair of vertices *U*_3,*j*_ and *V*_3,*j*_ for the variable setting path for literal *z*_3_. The root of the species clause gadget for clause *C*_*j*_ is a vertex *U*_*j*_ with *U*_1,*j*_ as one child and 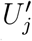 as the second child whose children are *U*_2,*j*_ and *U*_3,*j*_. Note that *U*_1,*j*_, *U*_2,*j*_, and *U*_3,*j*_ are hybridization nodes since they each have two parents. Node *V*_1,*j*_ has a second child *V*_*j*_, *V*_2,*j*_ and *V*_3,*j*_ have a child 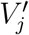 (a hybridization node), 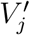 is another parent of *V*_*j*_ (a hybridization node) which, in turn, has a single child *𝓁*_*j*_, a leaf of the species tree. The roots of the *m* species clause gadgets are connected with a perfect binary tree *S*^*c*^.

Finally, the species variable gadget tree *S*^*x*^ and the clause gadget tree *S*^*c*^ are connected using two 2*n*-combs called *D* and *E* analogous to the *d* and *e* 2*n*-combs in the gene tree. Specifically, the root of the species tree has two children, one of which is the root of comb *D* and one is the root of comb *E*. One of the two deepest leaves of comb *D* is the root of the variable gadget tree *S*^*x*^ and the remaining leaves are labeled *D*_1_, …, *D*_2*n*_ in increasing depth from the root. Similarly, one of the two deepest leaves of comb *E* is the root of the variable gadget tree *S*^*x*^ and the remaining leaves are labeled *E*_1_, …, *E*_2*n*_ in increasing depth from the root. A representation of the species network is shown in Figure 2.

**Figure 2.**
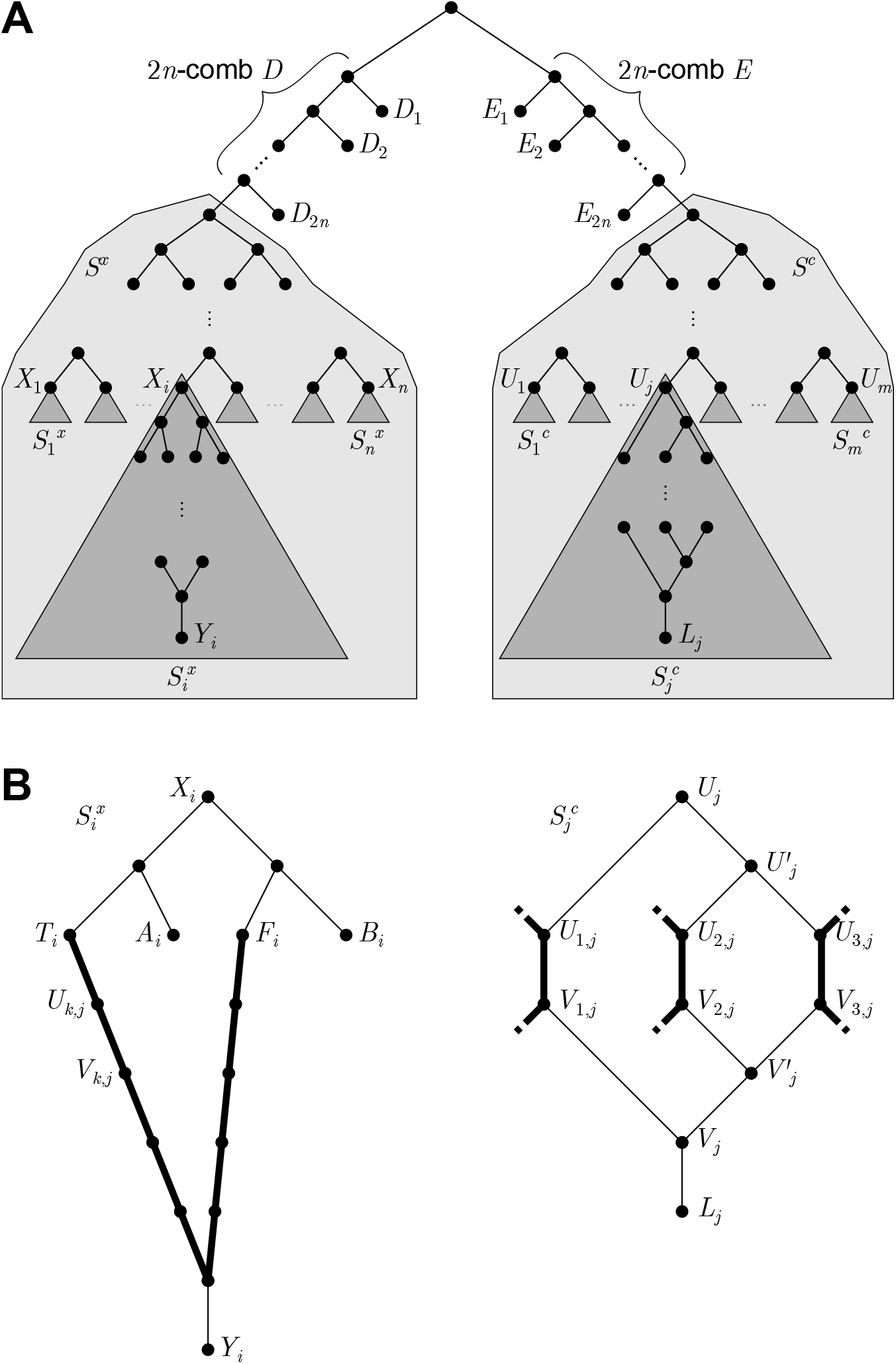
The species network in the NP-hardness reduction. (A) The shaded subtree *S*^*x*^ on the left contains a variable gadget 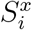 for each variable *x*_*i*_ in the 3SAT instance, and the subtree *S*^*c*^ on the left contains a clause gadget 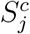 for each clause *j* in the 3SAT instance. (B) The variable gadget on the left is shown in detail. Vertices *T*_*i*_ and *F*_*i*_ are the first vertices on the variable setting paths (indicated in bold) for variable *x*_*i*_. The clause gadget on the right is shown in detail for clause *j*. The bold edges are from variable setting paths for the three variables in clause *j*. Note that the edge *U*_*k,j*_, *V*_*k,j*_ indicated on the right is the *k*^*th*^ bold edge in the clause gadget for clause *j* iff variable *x*_*i*_ is the *k*^*th*^ variable in clause *j*. In this example, *U*_*k,j*_, *V*_*k,j*_ appears on a true variable setting path, indicating that variable *x*_*i*_ appears unnegated in clause *j*.

The leaf map *Le* is as follows: For the leaves in the variable gadgets, *Le*(*a*_*i*_) = *A*_*i*_, *Le*(*b*_*i*_) = *B*_*i*_, and *Le*(*y*_*i*_) = *Y*_*i*_, for 1 *≤ i ≤ n*. For the leaves of the 2*n*-comb gadgets, *Le*(*d*_*i*_) = *D*_*i*_ and *Le*(*e*_*i*_) = *E*_*i*_, for 1 *≤ i ≤* 2*n*. For the leaves in the clause gadgets, *Le*(*𝓁*_*i*_) = *L*_*i*_, for 1*≤ i ≤m*.

Finally, the value of *k* in the decision problem is set to be *n*, the number of variables in the 3SAT instance. It is easily seen that this construction can be performed in time polynomial in the size of the 3SAT instance.

### Correctness

We prove that the constructed MPRD instance has a reconciliation with deep coalescence cost no more than *n* if and only if there is a satisfying assignment of the variables in the given 3SAT instance.

We begin with several observations about any lineage map *M* corresponding to a reconciliation for the constructed instance of MPRD. The species network *S* has unique paths from its root to the leaves *A*_1_, …, *A*_*n*_, *B*_1_, …, *B*_*n*_, *D*_1_, …, *D*_2*n*_, and *E*_1_, …, *E*_2*n*_. Therefore, there-fore, all lineage maps share the same unique paths *M* (*𝓁*) for all leaves *𝓁* among *a*_1_, …, *a*_*n*_, *b*_1_, …, *b*_*n*_, *d*_1_, …, *d*_2*n*_, and *e*_1_, …, *e*_2*n*_. The only leaves *𝓁 ∈ L*(*G*) for which *M* (*𝓁*) has more than one possible path are *y*_1_, …, *y*_*n*_ and *𝓁*_1_, …, *𝓁*_*m*_.

Note that in order for a reconciliation to have cost no more than *n*, the induced lineage map *M* must satisfy the properties that *M* (*y*_*i*_) contains the node *X*_*i*_, the root of the gadget 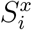. Similarly, *M* (*𝓁*_*j*_) contains the node *U*_*j*_, the root of the gadget 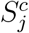. To see this, suppose by way of contradiction that there is a variable leaf *y*_*i*_ such that the path *M* (*y*_*i*_) (a path from *r*(*S*) to *Y*_*i*_) does not contain the vertex *X*_*i*_. The only paths from *r*(*S*) to *Y*_*i*_ that do not contain *X*_*i*_ are through clause gadgets, and therefore *M* (*y*_*i*_) must pass through the *E* comb. Since there is a unique path from *r*(*S*) to *A*_*i*_ in *S* and that path does not pass through the *E* comb, *M* (*a*_*i*_) cannot contain nodes from the *E* comb. Therefore, *M* (*y*_*i*_) must diverge from *M* (*a*_*i*_) at *r*(*S*), and therefore also diverges from *M* (*e*_2*n*_) at *r*(*S*) since *e*_2*n*_ is more distantly related to *y*_*i*_ than *a*_*i*_ is to *y*_*i*_. But then each of the 2*n* internal nodes of the comb *E* has at least two lineages, one from *M* (*y*_*i*_) and one from *M* (*e*_2*n*_), contributing a cost of at least 2*n > n*, contradicting the assumed cost bound. By similar reasoning, *M* (*𝓁*_*j*_) must contain *U*_*j*_.

There are, therefore, only two possibilities for *M* (*y*_*i*_) – it either includes the variable setting path for *T*_*i*_ or the variable setting path for *F*_*i*_ in the variable gadget for *x*_*i*_. Both of these options contribute at least one to the total cost since *M* (*a*_*i*_) and *M* (*b*_*i*_) must diverge at (or above) *X*_*i*_ and, since *y*_*i*_ is more distantly related to *a*_*i*_ and *b*_*i*_ than *a*_*i*_ and *b*_*i*_ are to one another, *M* (*y*_*i*_) must diverge from *M* (*a*_*i*_) and *M* (*b*_*i*_) at (or above) *X*_*i*_. Thus, *M* (*y*_*i*_) must contribute an extra lineage on an edge shared with *M* (*a*_*i*_) or *M* (*b*_*i*_). Since there are *n* variables, this contributes a cost of *n*, so these are necessarily the only extra lineages. There are only three possible lineage maps for each *M* (*𝓁*_*j*_). Each one passes through the corresponding clause gadget 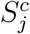, and therefore uses an edge on one of three species variable setting paths that pass through that gadget.

Now suppose there is a satisfying assignment of the variables in the 3SAT instance. Then construct a lineage map *M* with respect to *Le* as follows: *M* (*y*_*i*_) contains the variable setting path *T*_*i*_ if the variable *x*_*i*_ is set to true, and the variable setting path *F*_*i*_ if the variable *x*_*i*_ is set to false. For a clause *j*, let *z*_*k*_, *k ∈ {*1, 2, 3*}*, denote one of three literals in that clause that evaluates to true with respect to the given satisfying assignment. If *z*_*k*_ is an unnegated variable *x*_*i*_, then, by construction, the *F*_*i*_ variable setting path contains the edge (*U*_*k,j*_, *V*_*k,j*_) in the clause gadget 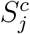. We then construct *M* (*𝓁*_*j*_) so that it passes through the clause gadget via that edge. If *z*_*k*_ is a negated variable *¬x*_*i*_, then, by construction, the *T*_*i*_ variable setting path contains the edge (*U*_*k,j*_, *V*_*k,j*_) in the clause gadget 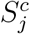 and *M* (*𝓁*_*j*_) is chosen to pass through the clause gadget via that edge. A reconciliation inducing this lineage map is trivial since each vertex in the gene tree has a corresponding vertex in the species network. The only cost incurred by this reconciliation is one for each variable gadget as noted above. The total cost is therefore *n* and thus this is a “yes” instance of MPRD.

Conversely, suppose there is some reconciliation with cost at most *n* and let *M* be the corresponding lineage map. Then, we induce a setting of each variable *x*_*i*_ based on whether *M* (*y*_*i*_) contains the *T*_*i*_ or *F*_*i*_ variable setting path. As noted previously, this induces a cost of *n* and thus the remaining paths cannot contribute any additional cost. Therefore, for each clause *C*_*j*_, *M* (*𝓁*_*j*_) must pass through an otherwise unused edge (*U*_*k,j*_, *V*_*k,j*_), *k ∈*{1, 2, 3}, implying that, by construction, the *k*^*th*^ literal in clause *C*_*j*_ has a setting that satisfies that clause. Therefore, the 3SAT instance is satisfied.

## 4 Discussion

In this work, we have shown that the problem of inferring an MPR between a gene tree and species network in the presence of incomplete lineage sorting is NP-hard. These results suggest several important directions for future research. First, approximation algorithms and exact fixed-parameter tractable algorithms should be explored for the MPR problem. Second, the problem may be solved effectively in many instances using satisfiability solvers or integer linear programming. Third, heuristics can be explored and tested experimentally.

## Acknowledgements

This work was supported by the Department of Computer Science and the Dean of Faculty of Harvey Mudd College. This material is based upon work supported by the National Science Foundation under Grant No.IIS-1751399 to YW and Grant No. IIS-1905885 to RLH. The authors thank Adam Walker for valuable comments that helped improve the paper.

## Footnotes

1 When explaining topological incongruence through only deep coalescence, a reconciliation is sometimes called a *coalescent history* (Elworth et al. 2019).

